# The enigmatic bone reduction pattern of thalidomide embryopathy is set by sequential exit from the sensitive window

**DOI:** 10.64898/2026.09.07.749812

**Authors:** Yossi Cohen

**Affiliations:** Adelson School of Medicine, Ariel University, Ariel, Israel; Institute of Hematology, Sanz Medical Center–Laniado Hospital, Netanya, Israel

## Abstract

Thalidomide reduces the limb bones in a puzzling order, starting with the thumb, the radius and the humerus and then returning to the ulna and the ulnar digits. To understand that order we analysed 435 thalidomide-affected limbs from the published record (1962–2025), of which 269 were upper limbs — 142 human and 127 non-human primate. Reconstructed by forward-chain assembly the reduction order runs: Digit 1 → Radius → Humerus → Digit 2 → Ulna → Digit 3 → Digit 4 → Digit 5. The co-reductions resolved into two rules. Within a segment (autopod, zeugopod), no bone is reduced before every bone anterior to it. Among the most posterior bones of the three segments (digit 5, ulna, humerus): no bone is reduced before every bone proximal to it. Behind these antero-posterior (A–P) and proximo-distal (P–D) reduction rules stands a maturation wave advancing postero-anteriorly and distoproximally. In developmental terms the bone precursors exit their sensitive windows after the autopod is built, in a biaxial order: from digit 5 towards digit 1, from the ulna to radius, and across segments from the autopod towards the stylopod (Digit 5 → Digit 4 → Digit 3 → Ulna → Digit 2 → Humerus → Radius → Digit 1). A co-reduction is therefore a snapshot of the prospective bones still sharing a sensitive window before their sequential biaxial maturation: exposure during elongation truncates the limb proximal to the progress zone, while later exposure reduces only the bones still immature at the time of exposure.

**Summary:** Thalidomide reduces the limb bones in a fixed order, attributed to sequential exit of each element from its sensitive window along two axes — antero-posteriorly within each segment (autopod, zeugopod) and proximo-distally among the three posterior elements (digit 5, ulna, humerus).

## Introduction

Thalidomide was introduced in 1957 and withdrawn in 1962 after causing more than 10,000 cases of severe congenital malformation worldwide (McBride, 1961; Lenz, 1962). A dose as small as 50 mg sufficed to produce embryopathy in humans, and four non-human-primate species proved comparably sensitive. The reduction order is stereotyped, running from the thumb to the radius, then the humerus, the index ray, the ulna, and last the remaining digits (Lenz and Knapp, 1962).

More than 30 hypotheses have been proposed, prominent among them neural-sclerotome damage, angiogenesis inhibition (Maier, 1965; Therapontos et al., 2009), oxidative stress (Parman et al., 1999), and the progress-zone model (Galloway et al., 2009). None recovered the order, nor explained why the humerus and the ulna, on opposite sides of the elbow, are lost at nearly the same point.

In 2014 I proposed that the stereotyped order reflects sequential maturation of bone elements postero-anteriorly, leaving the late-maturing preaxial elements susceptible to the insult for longer (Cohen, 2014). The model arose from a study by Koyama et al. (1996), who showed that cartilaginous chick limb elements acquire polarizing activity during their maturation in a posterior-to-anterior order — the inverse of the thalidomide reduction order. Here the reduction pattern was constructed step by step and then resolved into its components.

## Results

The cohort was assembled from 22 published human and non-human-primate sources and two online radiological repositories spanning 1962–2025 — the largest contributing series being those of Collins and Scott (2025), Hendrickx and colleagues (Hendrickx, 1973; Hendrickx and Newman, 1973; Hendrickx and Sawyer, 1978), Lenz and Knapp (1962), and Kida (1987), the last being the largest human series and the only one for which sixteen of its nineteen cases were checked limb by limb against the published radiographs and photographs; the remaining series contribute smaller numbers of element-resolved limbs, and the full source list, with per-source case counts, species and documentation, is given on the Source index sheet of Source data 1. Publications examined and not analysed, with the reason in each case and the number of rows each contributes to the record, are listed on the Excluded sources sheet of Source data 1. These yield 551 verbatim-coded upper-limb rows (355 human, 196 non-human-primate across four primate species), of which 355 record an affected limb; a further 80 affected lower limbs are recorded, 26 human and 54 primate, giving the 435 affected limbs of the corpus. For the analysis, the assembled data for each limb was reduced to the set of elements involved — thumb (T), radius (R), humerus (H), ulna (U) and digit rays 2–5 — giving 339 codeable affected limbs (210 human, 129 non-human-primate). Of these, 269 (142 human, 127 non-human-primate) have the eight elements assessable and so define an exact combination. The remaining seventy — sixty-eight human and two primate — are codeable element by element without reaching that standard. The limbs were divided into two groups before analysis. The truncation group holds those in which the loss is continuous from the stylopod through both zeugopod bones with the ulnar digits intact (n = 7 upper limbs, six human and one primate; and in the lower limb, the femur reduced, with the zeugopod taken whole or not at all, beneath an intact foot, n = 25, seven human and eighteen primate). The non-truncation group holds all others (n = 262 upper limbs, 136 human and 126 primate).

### Reduction order and rules

Reconstructed by forward-chain assembly from the co-reduced bones of the 142 human limbs, of which all but six are non-truncation, the reduction order runs: digit 1 → radius → humerus → digit 2 → ulna → digit 3 → digit 4 → digit 5. The arrows give the order in which the elements are reduced. Every adjacent step is unopposed in the 142-limb combination cohort. One further limb opposes one step: Lenz and Knapp 610315 right, element-resolved but without a complete combination because its radius is unassessed, reduces digit 2 while sparing the humerus, against sixteen limbs the other way. Its rays are scored from a finger count that names none of them — “Rechts drei Finger ohne Daumen” — so the identity of the second reduced ray follows from the preaxial reading rather than from the source. The order is also carried by the pairs that are not adjacent, and there it is stronger than the steps alone suggest — the eight elements make twenty-eight pairs, the 142 limbs supply 1,081 ordered observations across them, and 1,079 agree. The steps nonetheless rest on unequal numbers: 32 limbs discriminate the humerus from the radius and three digit 5 from digit 4 (Supplementary file 1). Within a segment, by contrast, the reduction order is fixed without exception (Autopod: digit 1 → digit 2 → digit 3 → digit 4 → digit 5; Zeugopod: radius → ulna), even when separated by elements of a different segment — two sequences interwoven into one. That is, no posterior element is reduced before all anterior elements of its own segment are reduced. This A–P reduction rule holds without a single exception across the 263 upper limbs (136 human, 127 primate) and the 80 affected lower limbs, from seventeen series and both species — 343 limbs in all, with no violation. In the lower limb it holds in both segments: within the autopod no digit is reduced unless every more preaxial digit is reduced, and within the zeugopod no limb reduces the fibula while the tibia — the preaxial bone, homologous to the radius — is spared, and none loses the fibula in isolation; in the macaque the tibia is reduced without the fibula in 13 limbs against none the other way. This rule implies an insult interrupting a specific developmental process. The same A–P polarity holds in thalidomide-exposed rabbits — the radius reduced while the ulna is spared, and the preaxial digit lost first — with no reported reversal in either segment (Pearn and Vickers, 1966; Vickers, 1967a, 1967b; Appendix 1).

Across segments the order is fixed among the three posterior elements, digit 5, the ulna and the humerus, and this is the P–D reduction rule: no distal element of the three is reduced unless every more proximal one is. In the human series and in Collins and Scott — the sources that document a limb bone by bone — it holds without exception, in 132 comparisons among 219 forelimbs and 16 among the lower limbs whose preaxial toe is reduced: the humerus reduced with the ulna spared in 49 limbs and never the reverse, with digit 5 spared in 66 and never the reverse, the ulna with digit 5 spared in 17 and never the reverse, and in the leg 3, 8 and 5 in the same three directions with none against.

When applied to the eight upper limb elements, the A–P reduction rule alone collapses the 255 combinatorially possible co-reductions to 35, and the two rules together to 25 (Table 1). Fifteen combinations are observed — nine in the human cohort and fourteen in the primate, and all but one of the human set falls within the primate one, the exception being 1-R-H-U. Fourteen of the fifteen are permitted. The fifteenth, 1-R-H-2-3-4-5, falls outside the proximo-distal rule; it rests on three primate limbs from one investigator’s alizarin-stained fetal material, occurs in no human limb, and is examined below. The eleven combinations the two rules permit that neither species shows are listed in Supplementary file 2. A third restriction on co-reductions, in addition to the antero-posterior and proximo-distal rules, is presented in the next section.

**Table 1.**
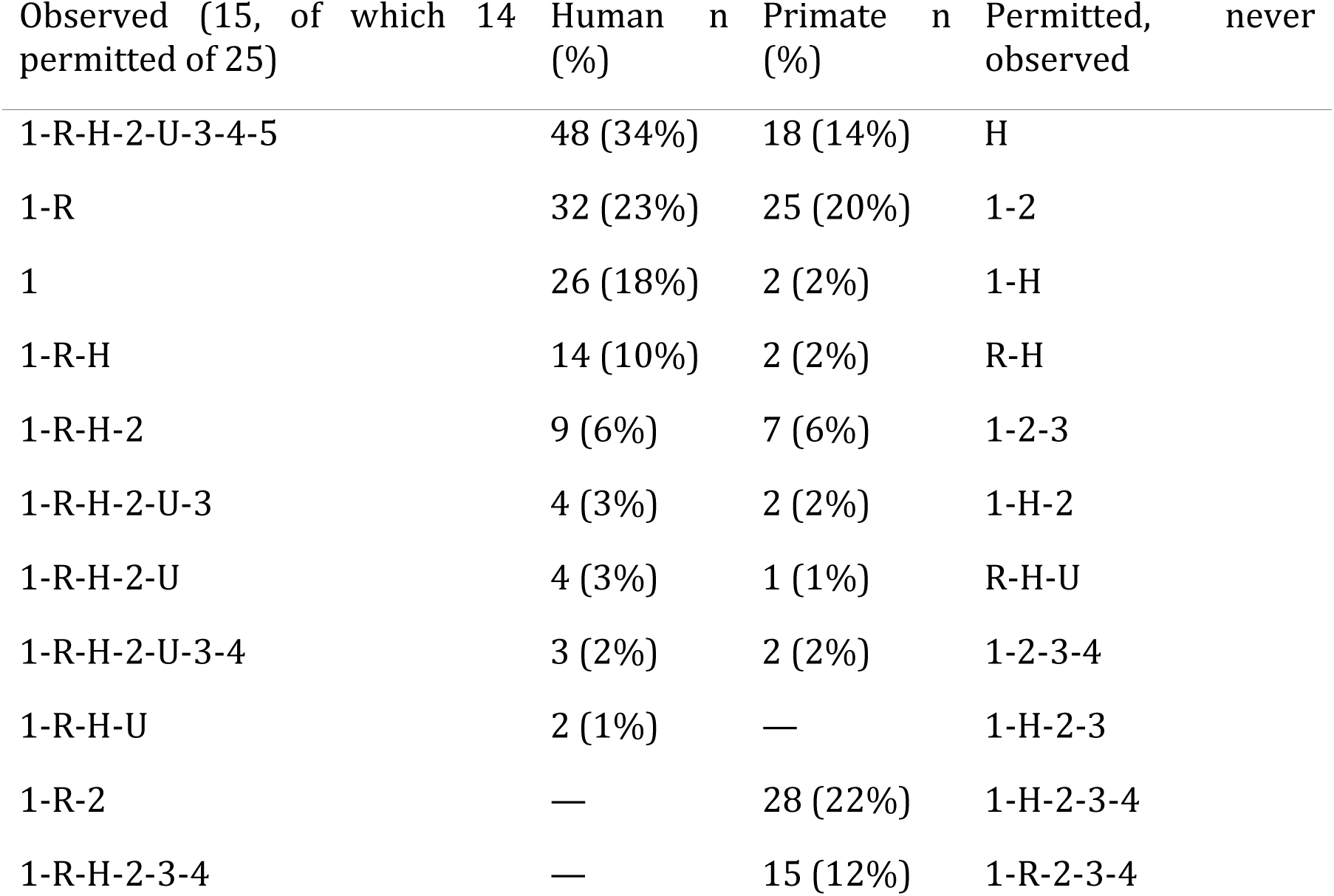
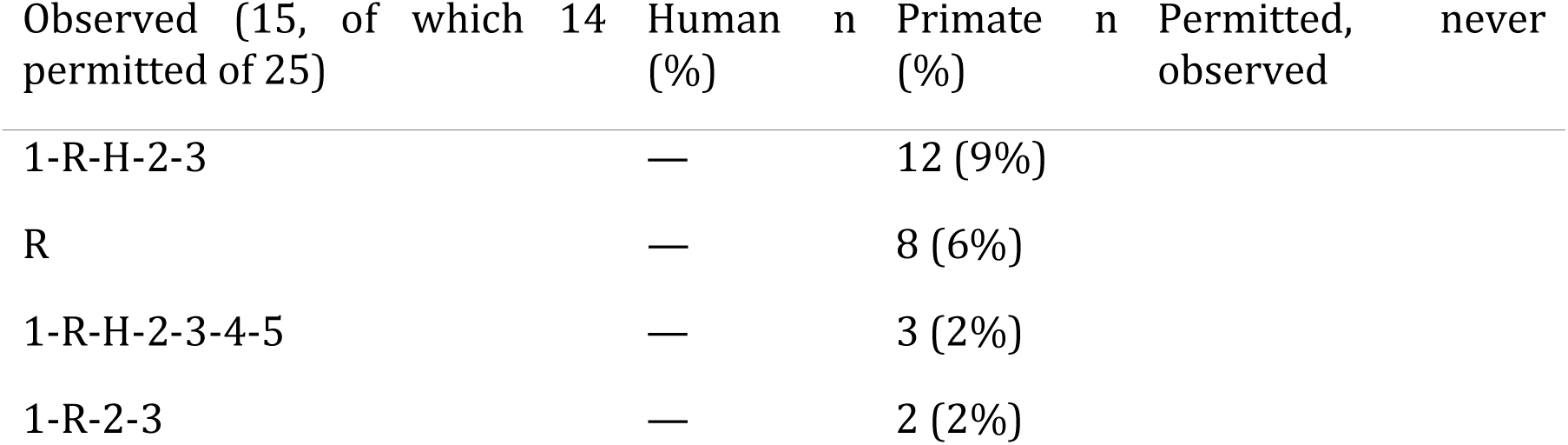
The co-reductions observed, and those the two rules permit but neither species shows. Each row of the first three columns is one combination of the eight elements, written as the elements reduced and separated by hyphens — 1 for digit 1, R for the radius, H for the humerus, U for the ulna, and 2 to 5 for the remaining digits, so that 1-R-H is a limb reducing digit 1, the radius and the humerus and no other element. Rows are ordered by the number of human limbs showing the combination; counts are limbs, with the percentage of that species’ cohort in parentheses (human n = 142, primate n = 127). An em dash marks a combination absent from that species. The fourth column is an independent list, not aligned row by row with the first three: it gives the eleven combinations the two rules permit that neither species shows.

### Shared sensitive windows

The hallmark of thalidomide embryopathy is the sensitive period in which the drug causes limb defects, about ten days. In terms of sensitive periods the duration of each element’s window is reflected by its reduction prevalence. An element-specific window is itself a conceptual shift, the sensitive period being usually considered a single interval shared by all elements. The A–P reduction rule is best attributed to sequential maturation of each segment postero-anteriorly. The P–D reduction rule is attributed to sequential cross-segment maturation from the autopod to the stylopod, so maturation advances in both directions (Supplementary file 3). As that model predicts, the reduction prevalence of the eight elements over the 111 element-resolved limbs of the series with a documented exposure window rises stepwise from digit 5 antero-posteriorly — digit 5 30% → digit 4 32% → digit 3 33% → digit 2 41% → digit 1 100% in the autopod, ulna 36% → radius 77% in the zeugopod, the humerus 54% alone in the stylopod — and along the postaxial column it runs from digit 5 through the ulna to the humerus, 30% → 36% → 54%. The three posterior rays lie within 3.5% of one another here and are resolved only in the Willert and Henkel series. The primate cohort gives the same directions, at ρ = +0.88 across the eight elements and +0.90 when the two cohorts are matched for depth by conditioning on the preaxial digit; the two rules alone force most of that agreement, so it is reported as consistency rather than as independent confirmation. Split by path the agreement is simpler still: the five digits and the three long bones each run in an order that is never broken, 520 ordered observations in the human cohort and 684 in the macaque with no violation in either. The two species differ only where the paths meet: the digits keep one order in both and so do the long bones, and what moves is where the two interleave, the ulna and the humerus standing later among the digits in man than in the macaque. Primate exposures were scheduled rather than random, so the same prevalences were also weighted equally by first dosing day — 124 of the 129 forelimbs carry one, spanning days 21 to 33 — and the weighting moves no element far enough to change the order, returning ρ = +0.88 again. Why the first dose is the one that counts is taken up in a later section. The sensitive windows are nested: the thumb’s window includes that of every other element, the radius’s includes the ulna’s, and the humerus’s includes those of the ulna and digit 5. The eight can therefore be set on a single scale of duration, starting from digit 5 and adding at each step the anterior and proximal elements that share a common window as postero-anterior and disto-proximal maturation advances. That is what Figure 1 shows for the human series, and Table 2 translates it into co-reductions accumulated from the first element to leave, digit 5, to the last two, the thumb at the end of the autopod path and the humerus at the end of the postaxial one.

**Figure 1.**
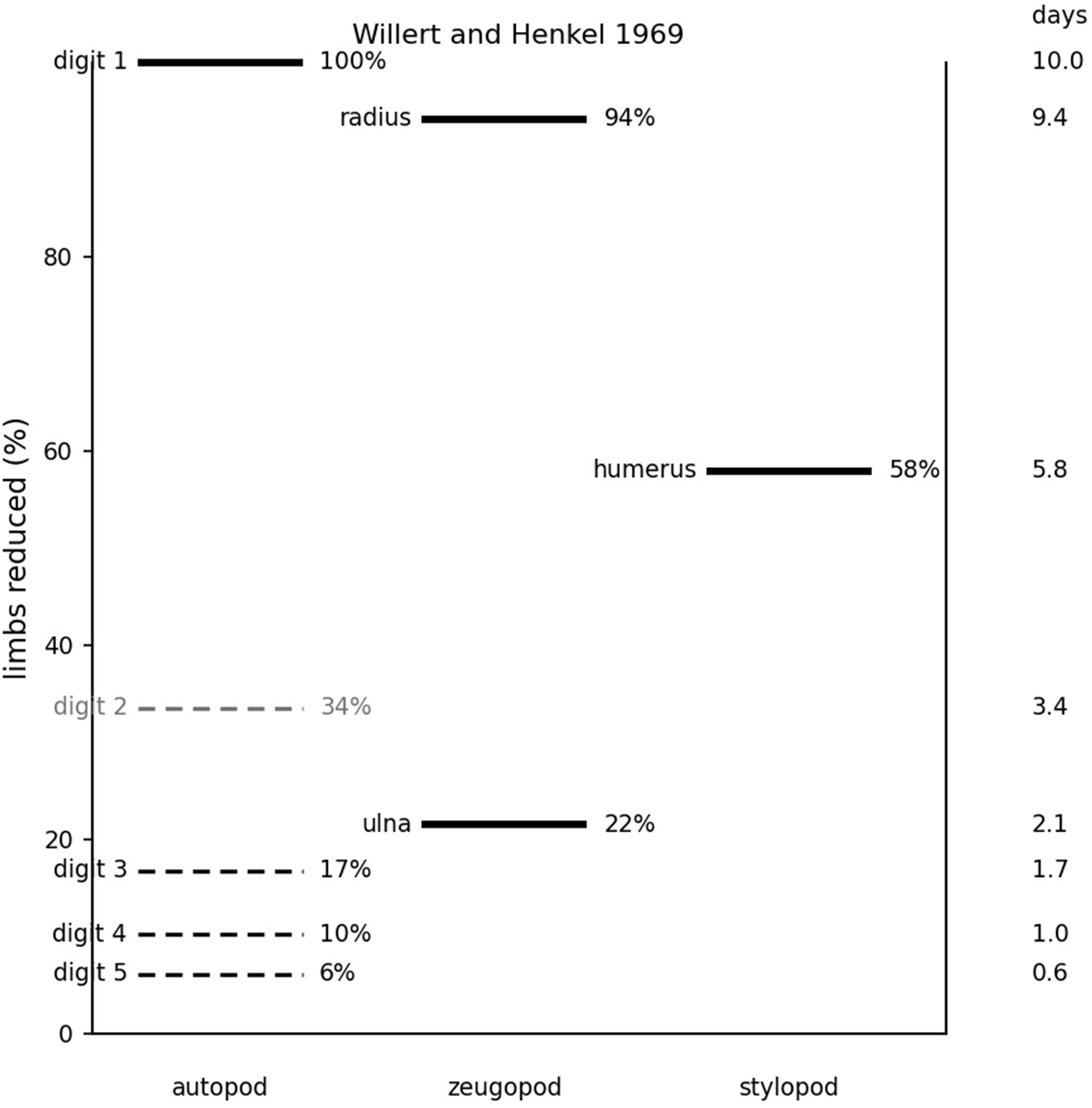
The sensitive windows of the upper-limb skeleton. Drawn from the 311 reduced upper limbs of Willert and Henkel (1969), the one series large enough and unselected enough to separate digits 3, 4 and 5. Each element sits at its reduction prevalence, read as the proportion of exposures that found it still vulnerable. Solid lines are counted from the sub-type tables; dashed lines are derived from published mean finger numbers and carry no combination for any individual limb. Digit 2 is grey, being the one element whose rank the data do not fix. The ulna is scored as in the cohort — a hypoplastic ulna counted only where an ulnar-side digit has gone — and comes to 21.5%; read at face value it would be 33.4% and stand level with digit 2. The vertical spacing ranks the elements and does not imply that every adjacent pair is resolved: the separation of digits 3, 4 and 5 here is drawn from this one series, and in the pooled prevalence cohort those three lie within 3.6 percentage points of one another, and within 6.3 when the ulna is included and are ordered by the co-reductions rather than by prevalence. The right-hand scale re-expresses prevalence as sensitive-period duration, closing on digit 1 at the accepted ten days: a proportional re-expression, not an independent estimate. The macaque series is reported as a rank correlation in the Results and is not drawn here, its exposures having been scheduled on chosen days rather than falling at random. Sourcing, the derivation of the dashed lines and the per-element values are given in Supplementary file 3. Willert and Henkel print every radiograph as a left extremity irrespective of the true side, for ease of comparison (p. 14 footnote), so the side recorded for those limbs is the side as figured; only seven of their rows carry a verbatim that names a side at all, and 53 of the 67 are coded left. Nothing in the analysis turns on side, since a combination is a set of elements, but bilateral statements exclude that series.

**Table 2.** The human exit sequence. Each row is the limb after one further element has left its window, so a limb exposed at that step reduces the elements still listed; n = 142.

| **Step** | **Element that has exited** | **Along** | **Still reducible** | **Limbs (%)** |
| --- | --- | --- | --- | --- |
| 1 | — |  | 1-R-H-2-U-3-4-5 | 48 (34%) |
| 2 | digit 5 | both paths begin here | 1-R-H-2-U-3-4 | 3 (2%) |
| 3 | digit 4 | antero-posterior | 1-R-H-2-U-3 | 4 (3%) |
| 4 | digit 3 | antero-posterior | 1-R-H-2-U | 4 (3%) |
| 5 | ulna | proximo-distal | 1-R-H-2 | 9 (6%) |
| 6 | digit 2 | antero-posterior | 1-R-H | 14 (10%) |
| 7 | humerus | proximo-distal | 1-R | 32 (23%) |
| 8 | radius | antero-posterior | 1 | 26 (18%) |

All but two of the 142 human limbs fall on this sequence. The exceptions are the two forelimbs of Kida 8012, which reduce the ulna while sparing digit 2 — a single patient, suggestive of a double hit at entry and exit (see the next section).

### Enter the sensitive period

In contrast to the classical embryopathy, which involves the autopod, an exposure that ends before the autopod exists should spare it. The sensitive period begins with the outgrowth, which is elongated by mesenchymal cells released by the progress zone, so the phenotype at entry is proximal truncation, starting from the stylopod. Proximal truncation is rarely documented in the upper limb, where one or more digits are usually missing as well. In the lower limb, however, truncation of the femur with or without tibia and fibula beneath a foot with no reduced element appeared in 1 of 26 affected human lower limbs and 14 of 54 primate (including Collins 6110, 6367, 6476). Yet, a second, later exposure can reach the distal limb the first exposure spared: the first exposure truncates, the progress zone then elongates the limb as far as the autopod, and a later exposure at exit reduces the preaxial elements. In the upper limb this combination accounts for seven limbs of both species, the six human ones coming from five children. Collins 4696, dosed on days 25 to 27, has the humerus and radius absent and the ulna reduced, with the hand standing beyond it and digits 1 and 2 reduced at exit. Kida 8012, whose mother took the drug daily, is the clearest human instance: humerus absent, radius absent, ulna reduced, and beyond that the carpus and four rays still standing, the thumb alone gone. The lower limb shows the same combination in both species, but the second event differs. In the primate it is a loss or a duplication of the hallux, in animals exposed over several days — reduction in 6449, duplication in 6346 and 6363. In the human it is a duplication only: six of the seven truncated limbs, from four patients, carry a doubled or bifid hallux or preaxial polydactyly, and none carries a preaxial reduction. When the ulnar digits are reduced as well, the exposure fell after the autopod was built but before the posterior rays had left their window — a single event early in the exit sequence rather than two. Because a limb reached at both entry and exit has by definition seen two exposures, the shared-window test does not apply to it. The A–P and P–D rules cut the 255 possible combinations to 25, and the shared window cuts those to the eight prefixes of the order; admitting entry and exit as two events raises that to ten, and every one of the 142 human limbs falls inside (Supplementary file 8). A unique occurrence, Collins animal 6438, has the femur, tibia, fibula and two preaxial toes reduced on the left, and on the right the femur reduced with toes 1 to 3 while the tibia and fibula are intact. This segmental skip of the right zeugopod is best explained by truncation of the femur at entry into the sensitive period together with reduction of the toes at exit. The drug does not harm the progress zone itself; only the precursor cells it has already released are susceptible. The sensitive period is therefore dynamic: it lengthens with the progress zone at the entry and shortens as postero-anterior maturation takes the elements out of it.

### Timing, not dose, sets the reduction

Experimental exposures separate the contributions of dose, schedule (repeated doses) and developmental stage. In the Collins and Scott macaque series (Collins and Scott, 2025), dose spans a sixteen-fold range (0.625–10 mg/kg) and contributes nothing. Holding the exposure days fixed at 26–28, the lowest dose reduces three and two elements and the highest reduces two and two; at days 27–29 an animal given 2.5 mg/kg (eight and six elements) is more severely affected than several given four times as much. The per-limb response is all-or-none rather than graded.

Repeated dosing adds nothing at the exit. Holding the first dose at day 27 and the dose at 10 mg/kg, one day, three days and ten days give the same combination; the three animals given a single dose, on days 25, 27 and 29, are affected in all six forelimbs; and an animal dosed for ten days from day 28 is entirely normal in both. The hindlimb differs only in that its window is easier to miss. All six hindlimbs of the ten-day courses lose a long bone, against 17 of the 68 from three-day courses; but no ten-day limb loses more than the deepest three-day limb, three bones in each. A longer course raises the chance of catching the window and not the depth of what it takes, which is what a single hit predicts. The dose result carries the same meaning by a second route: because elimination is first-order, a higher dose necessarily prolongs the time spent above any effective concentration, so a sixteen-fold dose is itself a substantially extended exposure. Neither more drug nor more days alters which elements are lost. The per-animal working is given in Supplementary file 6.

The reduction order is the decisive evidence. Were the phenotype the sum of what each dose destroyed on its own day, an exposure spanning several days would superimpose the sets vulnerable on each, losing postaxial elements while intermediate ones survived and so producing patterns that violate the A–P reduction rule. No such limb exists: all seventy-six affected Collins forelimbs obey the rule, and so do all 142 human limbs. The limb responds as though it encountered a single developmental stage, not an accumulation of daily insults.

The dosing calendar makes the same point independently of prevalence. In the macaque series, whose gestational clock is common, the ulna ceases to be reducible after day 27 while digit 1, the radius, the humerus and digit 2 remain reducible to the last day of the series — the posterior element exits first and the anterior ones last, read from the experimental calendar and not from any frequency. The calendar also bounds the window: in the cynomolgus and bonnet series the forelimb window is days 25 to 28, four days wide, with a hard edge at each end. In the hindlimb the entry phenotype appears on days 27 and 28 and on no other, and the exit phenotype on days 28, 30 and 32 — two days against five, so the limb reaches its autopod about twice as fast as it matures. The rhesus series looks wider than this and is not. Every forelimb whose course begins on day 21, 22 or 24 is reduced, and so is every one beginning on day 33, which read literally would make the window twelve days; but no animal in that series received a single early dose — each early course runs three days — so a course beginning on day 21 and one beginning on day 27 cover much of the same ground, and the first-dose day indexes a course rather than a moment. Read across the whole series the day carries little information: depth falls only weakly with it (Spearman rho = −0.27 over 74 forelimbs), and courses beginning at days 21–26 and at days 27–29 are almost indistinguishable, digit 1 reduced in 33 of 34 against 33 of 36. Where single doses are available the edge is sharp instead: one dose on day 23 and one on day 24 gave no defect at all, while one on day 27 gave phocomelia of all four limbs (Wilson and Gavan, 1967). A four-day window with a hard edge, sampled by three- to ten-day courses, is what both observations give. The per-day counts are given in Supplementary file 6.

One series lets the two windows be watched sliding past each other. Collins and Scott dissected every animal and scored all four limbs whatever their state, so the forelimb and the hindlimb of one side can be read together in 74 animal-sides. Taking how much of the forelimb is reduced as a proxy for where in its window the exposure fell, the hindlimb does not worsen with it but rises and falls: with one to three forelimb elements reduced the leg loses 0.17 of its three long bones on average, with four or five it loses 1.25, and with six to eight it falls back to 0.50 (permutation p = 0.0001 for the rise and 0.07 for the fall, both limbs fully assessed). Five further animal-sides carry an untouched forelimb beside a reduced leg — 5703, 6476 on both sides and 6438 on both, the last losing all three leg bones with the arm entire. The sequence runs leg untouched, then leg worst, then leg recovering, then leg alone, and it is what two windows offset by a few days must produce: an exposure early enough to catch the arm entering its window finds the leg still a bud with nothing to lose; one in the middle finds both limbs open at once; one late enough to find the arm deep in its exit finds the leg’s proximal bones already mature; and one later still misses the arm altogether. The five phases are not separate phenomena but one pair of windows passing. Either end of the hindlimb calendar is empty: limbs first dosed on day 25 are unaffected, ten of ten, the leg being still a bud with no element to lose, and by day 41 nothing is affected again. The stage in which the leg is stylopod and zeugopod without a foot is a day or two wide, which is why pure phocomelia is the phenotype one catches only by dosing into it: fourteen hindlimbs in the whole cohort against nineteen with both ends involved. It also explains a contrast that has nothing to do with the leg being more vulnerable than the arm. Classical phocomelia is 14 of 68 affected hindlimbs and 2 of 347 forelimbs, and the reason is the schedule: these exposures were timed at the forelimb, and a dose aimed at the arm’s exit falls on the leg’s entry. These limbs, and every other the model has to account for individually, are set out case by case in Supplementary file 4.

## Discussion

The current analysis rests on a single body of data, the co-reductions recorded over six decades.

In summary, the reductions of 435 affected limbs were assembled from 22 published sources (1962-2025), coded element by element, and the co-reductions of the 269 that define an exact combination gave the reduction order by forward-chain assembly: Digit 1 → Radius → Humerus → Digit 2 → Ulna → Digit 3 → Digit 4 → Digit 5. A deeper assessment resolved this order into three interwoven sequences. Two run antero-posteriorly within a segment (Autopod: Digit 1 → Digit 2 → Digit 3 → Digit 4 → Digit 5; Zeugopod: Radius → Ulna), and both hold without exception, confirming the postero-anterior maturation proposed (Cohen, 2014). The third runs proximodistally among the most postaxial elements: Humerus → Ulna → Digit 5. In the sources that document a limb bone by bone it holds without exception in both limbs, across human, cynomolgus, rhesus and bonnet macaque and green monkey material, and under every definition of reduction tested. Read through prevalence as a surrogate for sensitive window duration, the ranking of the eight elements recovers on its own the same sequence that forward-chain assembly constructed from the co-reductions observed, and both the forward chain and the prevalence ranks agree with the same reduction order: Digit 1 → Radius → Humerus → Digit 2 → Ulna → Digit 3 → Digit 4 → Digit 5. The reduction order starts from digit 1 — that is, at the exit from the sensitive period, which is the inverse of the maturation order: Digit 5 → Digit 4 → Digit 3 → Ulna → Digit 2 → Humerus → Radius → Digit 1. However, exposure preceding the autopod’s completion truncates the limb proximally; truncation with the autopod wholly intact was recorded in this cohort only in the lower limb, while in the upper limb the seven truncated limbs all carry a preaxial loss from a presumed second exposure at exit, a double hit.

Thalidomide is a maturation probe of the limb bone precursors. That places the sensitive period entirely before ossification, which begins later and proceeds proximo-distally, so the sensitive window ends still on the cartilaginous model. Collins 6449 shows the probe reading four limbs of one animal at four positions in the sequence, and the positions differ by single steps. In the forelimb the right takes the humerus, the radius and digits 1 to 3, and the left takes those and the ulna; in the hindlimb the right takes the femur, the tibia and the hallux, and the left takes those and toe 2. The left is one element deeper on both limbs, and within the elements the right is caught earlier still — its tibia is four-fifths reduced where the left has lost it entirely, and its metatarsals stand where the left has lost the first two. One exposure, four limbs, and the differences between them are the next step of the order in each.

The insult itself is a single hit at the exit, timing-dependent and dose-independent, all or none. At the entry the proximal truncation follows the progress zone, increasing up to the end of elongation. At the exit the co-reductions decrease from amelia to preaxial reductions alone. Exposure on days 28 to 30 (Collins 6110) leaves the upper limbs at their preaxial exit, the radius and digits 1 and 2 reduced, while the hindlimbs are still at their entry with the femora absent beneath intact feet: the two stages fall inside a single three-day course, so the lag between the exit of the upper limbs and the entry of the lower is no more than three days. Collins 6363, dosed on days 27 to 29 at four times the dose, is one step earlier in both limbs — the arms losing the radius, with digit 1 on the left only, and the femora reduced at their distal fifth rather than lost. Collins 5350, dosed on days 26 to 28, has all four limbs at their exits, the arms losing the radius with digits 1 and 2 and the feet losing the preaxial toes — the same offset, one stage on. Nominal dosing days cannot order these animals against one another, since gestation is not fixed to a day, but each animal dates itself.

Thalidomide embryopathy exhibits further enigmas, among which extra bones — a triphalangeal and/or a duplicated ray — coexisting with reductions in the same limb. The Collins series bears on that: the supernumerary elements are themselves preaxial and are accompanied by preaxial reductions, in all five animals that carry one (4595, 5703, 6346, 6349, 6363), and in no affected limb is a duplication postaxial. That suggests that an impending insult to the preaxial ray stimulates a compensatory response by the nearby progress zone. Why the preaxial ray and no other? Because it has enough time to recruit fresh precursors from the adjacent progress zone: it stays a precursor longest, and is the element most often reached in either limb. Digit 1 is reduced in 97% of affected hands where digit 2 is reduced in 57%, and toe 1 in 47% of affected feet where toe 2 is reduced in 35%. Duplications are commoner in the foot — 24 of the 80 affected lower limbs carry one against 13 of the 355 affected upper — presumably because the greater distance the progress zone must cover in the leg leaves each element longer as a precursor. The one postaxial duplication in the material, a doubled fifth toe in both feet of Lenz and Knapp 610123, occurs in legs the source records no bone reduction in, and so is not a response to an impending loss and not necessarily a thalidomide insult.

Two predictions were made from the maturation sequence model before the data that test them were assembled, and they are independent of each other. The first is that reduction prevalence stands for the duration of an element’s sensitive window, so that the prevalence order and the reduction order must coincide. In the human cohort they do exactly, and by two different routes: the order was assembled from within-limb co-reductions, the ranking comes from marginal counts, and the two could have disagreed. The primate prevalences cannot serve the same purpose: their exposures were scheduled rather than random, as an estimate of duration requires, and reweighting by first dosing day corrects that only in part. The second is that the sensitive window comprises two processes advancing along two axes rather than a fixed interval, so that severity must rise and then fall as exposure moves through it; in 74 animal-sides the hindlimb loss climbs from 0.17 of three long bones to 1.25 as the forelimb deepens and falls back to 0.50 (p = 0.0001 for the rise). Neither was fitted to the material: the first follows from the 2014 proposal that maturation timing sets the pattern, the second from the proposal itself.

### Limitations

The three routes to the order are not independent bodies of evidence: all three read the same limbs, and their agreement is coherence rather than replication. What separates them is that a ranking cannot be violated while a rule can, so only the rules were exposed to refutation. The order is left open where the rules are silent, every departure in either species lying in the cross-path set, a long bone against a digit, which no rule here orders. And the coding rests on published descriptions: where no plate, radiograph or illustration accompanies a case, the verbatim is open to interpretation.

Two conventions bear on the counts. An element a source does not name is counted as normal, so a bone reduced but unreported is scored as spared and every prevalence is a lower bound; the Hendrickx and Newman material shows the same fetus recorded in Table 2 as a bilateral meromelia and illustrated in Figure 1C with the left forelimb absent, so the omission can be substantial. Against this, their skeletons were cleared and stained with alizarin red, which shows mineralised bone only: the authors report the carpals unossified at examination, the digits ossify later than the long bones, and the series mixes fetuses examined at 70 and at 100 days without stating which entry is which. A ray recorded as absent in that source may therefore be a ray not yet ossified, and its digit prevalences are correspondingly upper bounds. Neither convention affects the two rules, which compare elements within a limb rather than counts across limbs, and neither reaches the long bones: the radius and ulna ossify together, so the radius-absent-with-ulna-present pattern in all nineteen affected forelimbs of that series is anatomical. The remaining strengths and limitations — the counts the two rules permit, the four limbs that fall outside the P– D reduction rule, the conditioning of the lower limb, and the coding account — are set out in Supplementary file 7.

### Future directions

Several factors hold limb precursors undifferentiated: FGF, sonic hedgehog and Gremlin1, by antagonising BMP, keep that pool labile; collapse of the loop would release everything still uncondensed at once (Verheyden and Sun, 2008; Bénazet et al., 2009). Retinoic acid and other proximal signals drive maturation instead, which would free the humerus first, the opposite of the observed order. None of these factors, by expression or by ending, accounts for maturation that runs anteriorly within a segment and proximally across segments. SALL4, degraded when thalidomide binds cereblon (Ito et al., 2010; Donovan et al., 2018; Matyskiela et al., 2018) and switched off across the period (Akiyama et al., 2015), fits competence rather than a clock. Posterior digits leave their windows at nearly the same time, yet the antero-posterior order is never broken, suggesting that the maturation front advances anatomically in a continuous manner postero-anteriorly and crosses the whole mesenchymal thickness to fixate the entire structure. This possibility is supported by the insult to limb connective tissue in addition to bone, with the same preaxial bias: twenty upper limbs in this cohort record an insult to the thenar eminence and none to the hypothenar, fourteen of them also reducing a skeletal element. The thumb points the same way: four patients here carry a triphalangeal thumb with no element reduced, so at that moment the ray was still open to being built as a finger — an effect on what an element becomes, not on whether it forms. A global switch from an embryonic to a mature expression signature has been described in brain, occurring over about two days in the neonatal mouse (Cohen, 2024). Whether the exit of limb elements from their sensitive window represents a comparable mesenchymal fixation, running on a different spatiotemporal schedule from patterning, identity and ossification, remains to be tested. The nature of the insult is also open: part of what is lost may be cartilage that formed and then failed, and never ossified, which would raise the possibility of a false commitment of the mesenchymal precursors.

## Materials and methods

### Cohort assembly and coding

A unified cohort of 551 verbatim-coded thalidomide-affected upper-limb rows was assembled from 22 published sources and two online radiological repositories (355 human, 196 non-human-primate: cynomolgus, rhesus and bonnet macaque, green monkey); the sources are listed in full, with per-source case counts, species and documentation type, on the Source index sheet of Source data 1. Inclusion required documented maternal exposure during the sensitive window, limb-by-limb radiographic/photographic/anatomical/3D-CT documentation, and absence of an alternative diagnosis. A case whose own source denies a thalidomide aetiology was never eligible, and such cases are held on the Excluded controls sheet of Source data 1 rather than entering the cohort and being removed from it: Coodin Cases 2 and 3 (1962), the two persons of Smithells and Newman’s Figure 5 (1992), and the three Zurich autopsy cases of Willert and Henkel (1969, pp. 117–118), which those authors present as phenocopies and of which they write that a thalidomide aetiology is out of the question, the children having been born years after the drug was withdrawn in Switzerland. Sixteen limb-rows are held out on this ground.

Each of fourteen upper-limb elements (humerus, radius, ulna, carpals, MC1–MC5, Ph1–Ph5) was assigned one of five codes: A (absent), P (partial/truncated), H (hypoplastic), N (normal) or NA (not codable). Each limb was the unit of analysis. Eight rules governed what a source statement was taken to support, and were applied to the whole cohort rather than case by case.

1. Segment length names no bone. A statement describing the length of a segment rather than a bone - “Armverkürzung”, “kurze Unterarme”, “short forearms” - supports no code for any element within it, and both long bones were coded NA. Where a source names one bone and reports the segment as short, the shortening is attributed to the named bone and the unnamed bone is coded normal. Reading an unnamed element as normal is a convention and not an observation, and it carries most of the cohort: 310 of the 330 long-bone codes of normality rest on a source not naming the bone rather than on a statement that it was intact. Loss of the radius alone shortens the segment: the ulna bows, the carpus displaces radially, and the forearm measures short although the ulna itself is of normal length, a secondary relationship established in the original descriptions (Willert and Henkel, 1969). In this cohort the ulna is intact in every limb in which the radius is absent and imaging resolves both bones. Ten upper limbs and two lower limbs are coded on the first basis and six upper limbs on the second. No separate rule is therefore needed to decide when an inferred ulna counts as reduced, and none is applied. The consequence is a claim that can be checked directly: no limb in either species reduces the ulna while the radius is spared, and none reduces a digit while a more preaxial digit is spared — zero exceptions in 269 upper limbs from twenty-one sources. The five digit combinations observed are the five prefixes of the ray order; the remaining twenty-six of the thirty-one non-empty subsets do not occur.
2. A soft-tissue or joint finding is not a reduction. Contracture, cutaneous webbing, syndactyly and clubhand describe digits that are present; none states that a phalanx is short, absent or truncated, and the elements they name were coded N or NA accordingly.
3. A clause naming a side applies to that side only. Where a record carries a single description for both limbs, a clause marked left or right was read as evidence for that limb alone; the contralateral element is unstated, not normal, and was coded NA.
4. Where a case has both a radiograph and a caption, the film is the element-level evidence. A caption may name more elements than the image resolves; the image governs.
5. Patterning changes are not reductions. A triphalangeal thumb is a supernumerary phalanx, not a missing one, and was coded N at Ph1 and recorded on a separate flag, as were fusion and preaxial duplication.
6. NA is not N. NA marks an element the source does not assess and was excluded element-by-element from every denominator. A diagnostic term was never read as an element-level observation: Lenz and Knapp (1962) define classical phocomelia by the presence of all five fingers and state that this form does not occur in thalidomide embryopathy, so “Phokomelie der Arme” reports severe proximal reduction with the hand unspecified, and the digits of such records are coded NA. An element recorded as unossified rather than absent was likewise coded NA, since remnants may persist as cartilage invisible on an infant radiograph (Willert and Henkel, 1969, pp. 64, 126).
7. An equivocal ulna is counted only on corroboration. Rule 1 attributes the shortening of a segment to the bone the source names, so no ulna carries a code on a description that never named it. Such an ulna is counted as a genuine reduction only where an ulnar-side digit is also reduced, that being the co-reduction signature of unequivocal ulnar reductions in both species; where the ulnar digits are spared the shortening is treated as secondary to the radial defect and not counted. An ulna coded absent or partial always counts. Seven human limbs carry an equivocal ulna and the rule counts two of them. Its effect is reported in Supplementary file 5: without it the ulna rises from 36.5% to 40.9% and holds fifth place in the order either way, the limbs consistent with one exposure fall from 142 to 139, and the A–P reduction rule holds without exception either way. The reasoning is the authors’ own. Willert and Henkel describe the ulna of the transitional axial type as “vorhanden, aber meist deutlich verkürzt” (p. 35), and state that the ulna does not reduce by degrees as the humerus and radius do, being either present and clearly recognisable or wholly absent, its shortening elsewhere arising “wahrscheinlich erst als sekundäre Folge der Radiusreduktion” (p. 54). That the equivocal grade is the one that behaves differently can be shown by holding the radius constant. Among the limbs whose radius is wholly absent, an ulna coded absent or partial carries a reduced humerus in all eighty, a hypoplastic one in five of ten, and an intact ulna in eighty-four of a hundred and fifty-one. With the radial defect the same in every row the difference cannot be the radius: the hypoplastic grade behaves as an intact ulna does, and the two unequivocal grades do not. 8. A ray is reduced when its metacarpal or its proximal phalanx is lost. A ray that keeps both while losing phalanges beyond them is present and short, not missing, and the loss was recorded in the verbatim but not scored as a reduction of that ray. Seven rays in the material meet this criterion, found by searching every limb for one: digit 3 of a forelimb and the hallux or second toe of three hindlimbs in Collins and Scott, where the plates show the lost phalanges to be the distal ones, the verbatim giving their number but not which; the fifth ray of both forelimbs of one Hendrickx and Newman animal, where the source names them — “medial and distal” on the left, and on the right the medial phalanx alone with the distal one present, which is an intercalary loss within the ray rather than a truncation of its tip; and the third toe of one Hendrickx and Sawyer hindlimb, whose second phalanx alone is absent. The criterion is the metacarpal and the proximal phalanx and not the word terminal, so it covers both. All seven are coded alike. Scoring them as reduced instead would put one limb outside the A–P reduction rule and leave the other six unchanged, so the rule is stated on the coding that treats all seven alike rather than on the one that exempts none.

Records giving only a summary diagnosis (“phocomelia”, “meromelia”) or otherwise lacking element-level detail (135 upper limbs) were excluded from element-level analyses. Eighty-three of these limbs are described with the word phocomelia; in seventy-nine of them the digits had been assigned from the term and are coded NA. The cohort is limbs whose own source attributes the malformation to thalidomide taken within the sensitive period. A limb the source does not so describe is not excluded; it was never eligible, and the criterion it fails is recorded on its own row. No limb is set aside after the analysis: there is no post-hoc exclusion, and the tags are listed on the Inclusion criteria sheet.

Two publications were set aside in their entirety — Kowalski et al. (2015) and Tajima et al. (2016) — and are recorded with their reasons on the Excluded sources sheet of Source data 1, which also names the three that are coded and kept but contribute no analysed limb, among them Smithells and Newman (1992) and Malik and Cohen (2021): neither contains an element-resolvable limb, one reporting limb involvement only as a presence mark and the other yielding no limb with all elements assessable.

### Ineligible cases

No limb is set aside after the fact.

Every limb that does not enter the analysis fails one of the stated inclusion criteria, and the criterion it fails is named on its own row of Source data 1, so the funnel is reproducible from the data without reference to any list. A case whose source denies the aetiology is a control; a limb the source does not resolve element by element cannot define a combination; a limb the monograph illustrates twice is deduplicated against the figure legend that identifies the second view; a limb with no element reduced is not an affected limb. Two further criteria fall on one case each. Where one child appears twice in a source with findings that cannot both be true, as the birth date of 1 November 1961 does in two transcription blocks of Lenz and Knapp, neither record is usable and no limb of that child enters. Two macaque forelimbs (Collins and Scott, 2025, animal 5741) received D-thalidomide, the R-enantiomer, rather than the racemate, and were excluded by the original investigators on that ground; they are retained here, because thalidomide racemises in vivo with a half-life of the order of minutes, so an animal dosed with a single enantiomer is exposed to the racemate within the first hour and throughout a window traversed over days. The enantiomer administered is not the enantiomer the embryo meets. The decision is not load-bearing: withholding animal 5741 leaves the reduction order, the observed co-reductions and the zero A–P violations exactly as reported. Two human limbs (Ling et al., 1962, Case 2) had thalidomide documented from day 53 post-LMP, outside the 34–50 day sensitive period, together with a concurrent three-drug antiemetic that the reporting authors state cannot be excluded as a contributor; Case 1 of the same report, in which thalidomide was the only drug administered and from the third week, is retained. The phenotype agrees with the timing: that limb has the humerus absent with all five digits intact, and no limb of either species reduces the humerus while the thumb, the element with the longest window, is spared. The excluded pair is worth naming, because it is the one classical phocomelia in the material: both humeri absent, the radius and ulna present and fused to one another, and the metacarpals and fingers, in the reporting authors’ words, “formed normally”. That configuration — a reduced stylopod beneath an untouched hand — appears in none of the 269 limbs of the analysed cohort, and it appears here in a case set aside on exposure timing before its phenotype was considered. Two human limbs (Cuthbert and Speirs, 1963, Case 15) are reclassified as thrombocytopenia-absent radius syndrome rather than thalidomide embryopathy. Exposure is recorded as week 6-10 of pregnancy, beginning after the upper-limb window closes, whereas every other included case in that series begins on day 8-21 or is recorded as early pregnancy. The limbs show bilateral radial aplasia with the thumb preserved and digits 1 and 2 fused, and the source additionally reports an unusually long splenic flexure with adhesion of contiguous surfaces and absence of the appendix. Preserved thumbs with absent radii is the discriminating feature of that syndrome and the inverse of the thalidomide pattern, in which radius and thumb are lost together in every limb of this cohort; the visceral findings belong to the same syndrome. It was first delineated in 1969, six years after this series was published, and could not have been recognised by its authors. Platelet counts are not reported. The reclassification was decided on exposure timing and phenotype before the prevalence figures were recomputed. With these two limbs retained, digit 1 prevalence is 98% rather than 100% and every other element, the rank order, the single-exposure count and the zero A–P violations are unchanged. Two human hindlimbs (Hagen, 1965) lose the fibula with its muscles and their nerves, and the proximal femur with the acetabulum. They are not set aside but uncodable: Hagen never assesses the tibia, which appears once in the report and only as the landmark a fibrous strip is attached to, so it is NA under the rule that an element not assessed is not an element spared, and the limb cannot be scored for a rule that requires all three long bones assessable. No exclusion is load-bearing. The shallow combinations of the human series are not distinguishable, limb by limb, from non-thalidomide radial dysplasia. Willert and Henkel’s survey of the pre-1958 literature (1969, Tabelle 4) collects 169 cases of which 152 (90%) carry a radial aplasia or hypoplasia and 15 (9%) a thumb defect without one — the phenotype of thrombocytopenia-absent radius, Holt–Oram, Fanconi anaemia and the VACTERL association, one entry being labelled Fanconi syndrome by those authors themselves and three others explicitly familial. The combinations 1 (26 limbs) and 1-R (32 limbs) fall inside that spectrum, so 58 of the 142 human limbs are individually compatible with a background cause. What separates the series is not the frequency of any one pattern but its depth and its distribution: in the same pre-1958 material a humerus defect appears in 1% of cases, phocomelia in 2% and amelia in 2%, against a humerus reduced in 57% of the present human cohort, and roughly half the pre-1958 cases are unilateral where thalidomide dysmelia is characteristically bilateral with little side difference. Willert and Henkel set out five criteria on that ground — a documented ingestion, the 1958–1962 epidemic clustering, strict bilaterality, four-limb involvement and a clustering of severe defects — and every limb here is drawn from a source that applies them. The deep combinations, which carry the ordering, have no counterpart in the background series at all: it is the shallow limbs a background cause could explain, and the shallow limbs are not the ones the sequence rests on. No minimum occurrence count was applied, because one would cut in the wrong direction, retaining 1 and 1-R and removing the deep combinations that have no counterpart; applied at three occurrences it would also empty five of the macaque’s eight depths. Rarity here is a consequence of depth, not a mark of doubt. Reinstating every row on the sheet at once leaves the A–P reduction rule without an exception in either limb, the fibula still never reduced with the tibia spared, and never lost in isolation. No limb is excluded for the pattern it shows. Two macaque forelimbs (Hendrickx, 1973, day 27) report the humerus absent while both forearm bones persist, and were set aside in an earlier version of this analysis as anatomically unresolvable. They are retained: that was an argument from plausibility against a stated observation, and it fell on the only two limbs in the cohort in which the humerus is reduced while the radius is spared — the two forelimbs of one crab-eating macaque, dosed on day 27. Both satisfy the A–P reduction rule in each segment — the autopod is a proper prefix, the zeugopod untouched — so what they qualify is the cross-segment ordering, not the rule. They are also the two limbs that hold the primate single-exposure count at 115 while carrying the denominator from 125 to 127, both being inconsistent with any one exposure: on the primate prevalences one skips the radius and digit 2 at a cost of 21.3% and the other skips the radius at 1.6%, and no single exposure accommodates either. Phocomelia and amelia are both retained. In phocomelia the thumb, radius, humerus, digit 2 and ulna are reduced and the ulnar digits spared — an ordinary prefix of the reduction order, and the two- or three-digit limb of the published illustrations. Amelia is the complete co-reduction the rule permits and the sequence produces when exposure precedes every exit, and accounts for 48 of the 142 human and 18 of the 127 primate limbs. Amelia is not, however, simply the wave caught before any element had matured: in the primate series, where the first-dose day is known, the four limbs exposed earliest (days 21– 22) are all partial reductions and none is amelic, while the amelic limbs are distributed across days 24–28, the same days that mostly yield partial reductions. Amelia is therefore a minority outcome at ordinary exposure times rather than the endpoint of the sequence, and whether it belongs with the co-reductions at all is not settled by these data; we retain it because its elements are unambiguous, and note that excluding it would remove 49 human limbs and lower every prevalence without altering which combinations occur.

### Analysis subsets

Two analysis cohorts were used, each matched to its purpose. The reduction order, the A–P reduction rule and the co-reduction combinations (Table 1) were computed on all affected limbs with the eight ordered elements individually assessable, the digit rays being collapsed from their metacarpal and phalangeal codes, excluding only the cases that cannot be scored element by element and the few excluded on individual grounds, both set out above: 142 human and 127 non-human-primate limbs. This is the appropriate set for a combinatorial analysis, which depends on which elements co-occur, not on how representative the exposure timing is. Per-element reduction prevalence, interpreted as an estimate of sensitive-period duration, was instead restricted to six series from five authorships whose exposures sample the sensitive period broadly and near-randomly (Lenz and Knapp, 1962; Cuthbert and Speirs, 1963; Willert and Henkel, 1969; Kida, 1987; McCredie and Willert, 1999; n = 111). The choice of that cohort is tested in Appendix 3. Figure 1 is drawn from one of those series alone, because prevalence as a measure of vulnerable duration needs a denominator not selected on severity: amelia reduces all eight elements at once, so a series enriched for it lifts every line together and closes the intervals the figure exists to show. As published, the Willert and Henkel series is 5.9% amelic and Kida’s 6.9%, against 49% in Lenz and Knapp and 50% in Cuthbert and Speirs. The Lenz and Knapp material is published as two independently reported sets, the 1962/1969 series and the Figure 6b series, which are pooled here and held apart in the per-series reproducibility test, where splitting them can only make agreement harder to obtain (Supplementary file 1); series with few or timing-skewed cases — the small saturated single-case reports (Schuler-Faccini, Vianna, Kajii, Hagen, Ling, Coodin; 32 limbs) — were excluded from prevalence only and retained in the combination cohort, because prevalence requires a fair sample of exposure times that the combination analysis does not. The dose-calendar analysis, which reads the exposure day rather than the phenotype, is restricted further to the macaque series, whose forelimbs are reduced by exposures first given from day 21 to day 32 and whose hindlimbs from day 25 to day 33. Gestational day is not comparable across species whose organogenesis differs: in the green monkey it begins five to seven days later and the sensitive period runs from day 28, while Delahunt and Lassen (1964) dated gestation from the first day of menses and mated over an eight-day window, so that their day 32–42 exposures begin somewhere between the fifteenth and the twenty-second day after conception and cannot be placed on either axis; their own text sets that schedule against limb development “26 to 26.5 days after conception”, and, as a first dose preceding every exit predicts, three of their four forelimbs are amelic and the fourth retains a single digit. Both are set aside for that analysis alone and retained everywhere else, leaving 111 limbs with a recorded first-dose day; where an exposure spans several days the first is taken, the phenotype being fixed by the first dose. Last-reducible days were computed on the limbs whose elements can be scored individually, a wholly absent limb placing no element in time. Reduction was defined per cohort by documentation quality: inclusively in humans (absence, truncation or hypoplasia), as the clinical literature (1958–2025) does not reliably separate hypoplasia from absence, and strictly in non-human primates (absence or truncation), from their accurate experimental skeletal illustrations. The inclusive definition is applied consistently, so a femur recorded only as arched is a reduced femur, and a human limb whose only finding is hypoplasia of an element is a reduced limb: eleven such limbs enter the cohort on that basis, most of them the milder side of an asymmetric bilateral case whose contralateral limb is also analysed, and excluding them would remove from the denominator exactly those limbs in which most elements are spared. The bearing of that choice on the results is reported in Supplementary file 5: the A–P reduction rule holds without exception in both cohorts under every definition, and what the choice moves is how many limbs form an exact prefix of the order — 4 of 142 human limbs fall outside it under the inclusive reading against 18 of 120 under absence alone, because a limb whose intermediate rays are described as rudimentary or contractured is scored as sparing them when hypoplasia is not counted. Kida (1987) reports each patient three times over: a verbatim radiological description, an independent severity grade by segment, and acromion-to-fingertip limb lengths, alongside photographs and radiographs of most cases. None of the last three was used to derive a code. Used as checks they agree with the coding: the author’s own grade matches for all nineteen patients once his convention of grading joint involvement is translated to our convention of coding bone reduction, and the lengths match the coded laterality in four of the five patients for whom they are given, the fifth being a bilateral phocomelia in which the ordinal scale has reached its floor.

### Within-segment rule and prevalence

A limb violated the autopod rule if any digit ray was reduced while a more preaxial, assessable ray was spared (each ray one unit, reduced if its metacarpal or any phalanx was absent or truncated), and the zeugopod rule if the ulna was reduced while an assessable radius was spared. Per-element prevalence was the fraction of assessable affected limbs in which the element was reduced. Within-species reproducibility was tested within each randomly-timed series by an exact sign test on every adjacent pair of the pooled order, scored on the limbs of that series which separate the pair — one element reduced, the other spared — rather than by comparing derived orders, which at these sample sizes would read a sequence out of single limbs. Twenty-eight of the forty-two series-by-pair comparisons have limbs that separate the pair. Two run against the pooled direction: the humerus with digit 2, on a single limb, and digit 2 with the ulna, at one limb to five, which is also the pair the pooled data separate least. Six are individually significant in the pooled direction, in three different series (Supplementary file 1). Each prevalence was assigned an uncertainty by leaving out one series at a time; the resulting jackknife standard errors range from nought to seventeen percentage points, the lower bound being exact because the posterior rays carry identical prevalences and move together under every omission. Two adjacent pairs are separated by more than their uncertainty — digit 1 from the radius, and the radius from the humerus. Digit 3 and digit 4 coincide exactly and are not ordered by these data at all; the remaining pairs are separated by less than their uncertainty. The prevalences therefore fix most of the order but not the intervals between its steps. The per-element prevalences, read across the three limb segments, are tabulated in Supplementary file 3.

### Window duration and the shared rule

Soft-tissue findings were not coded as elements and enter no analysis; they are recorded in the verbatim and the coding note of the row they belong to, and the counts given in the Discussion are of rows whose verbatim names the structure. The thenar eminence is named on twenty upper-limb rows — 50, 165, 166, 175 to 178, 185 to 188, 195, 196, 199, 200, 203, 204, 396, 488 and 489 — and the hypothenar on none. Element coding was performed with the assistance of a large language model (Claude, Anthropic), which applied the coding rules set out here to the verbatim descriptions transcribed from each source. Where a source presents its findings as a skeletal illustration rather than as text — the alizarin-stained preparations of Collins and Scott (2025) — the illustrations were read by the author and rendered into verbatim text before coding. Every code was checked against the source by the author, who adjudicated each disputed case and recorded the decision, with its date and the source’s own words, in a changelog held with the data. The verbatim, the element codes and the coding note are deposited together for every limb, and every figure reported in this paper is regenerated from that workbook by the deposited scripts, whose verification harness checks each printed value against the data. Coding by one investigator and one model under one set of rules is a single reading of the record; no independent recoding exists, and the agreement between the two species tests the rules against different material but not against a different reader. A further limitation is diagnostic rather than editorial: SALL4-related disorders phenocopy thalidomide embryopathy closely enough that individuals reported as thalidomide cases have later been shown to carry SALL4 variants (Kohlhase et al., 2003; Kodytková et al., 2023). The human material here is drawn from a published record that predates molecular testing and may contain such cases. That bears directly on the one limb that departs from the reduction order: the element set reported for SALL4 loss — thumb, radius, humerus and ulna, with no reduction of digits 2 to 5 reported — is the combination Kida 8012 carries. Limbs were sorted into two categories before analysis. A limb was classed as truncation when the loss was continuous from the stylopod through both zeugopod bones — humerus, radius and ulna all reduced — while digits 3, 4 and 5 remained intact: an ulnar hand standing beyond a proximal loss is evidence that the autopod was built after the insult, so such a limb was reached at entry and again at exit and has by definition seen more than one exposure. Seven upper limbs meet this definition, six human and one primate, in two combinations (1-R-H-2-U in five, 1-R-H-U in two). The criterion is anatomical and was applied without regard to the cost a limb carries: five of the seven fall on the exit order at no cost at all. Its empirical warrant is the dosing record, where a loss continuous through both zeugopod bones never follows a single dose — none of the single-dose animals shows one, and the nine forelimbs of the dosed series in which the humerus, radius and ulna are all reduced were each given a course of three days or more — whereas a reduced humerus with the ulna spared, which is simply the antero-posterior order, follows a single dose readily. Within the group only a limb whose second event spared digit 2 records the two exposures distinguishably: where digits 1 and 2 have both gone the combination is an exact prefix of the exit order and is indistinguishable from a single deep exposure, so the configuration in which two events can be seen at all is narrow, and the cohort holds one instance of it. The category is correspondingly rare in the arm and common in the leg — seven upper limbs of six individuals against twenty-five lower limbs of thirteen — which follows from the arm’s window closing near day 24, before most exposures began, while the leg was still entering. All other limbs were classed as non-truncation. No limb is set aside on this basis. The whole cohort is scored together, and the category serves to interpret the limbs that depart from the reduction order rather than to remove them: a limb reached at entry and again at exit is expected to depart, and the combinations such a limb can take are enumerated in Supplementary file 8. A proximal truncation with the autopod wholly intact is a third configuration and occurs in the lower limb only. Each element’s reduction prevalence was computed in the fully-assessable human set and interpreted, under the single-wave model, as the fraction of exposure times at which postero-anterior maturation had not yet reached that element; a brief exposure reduces all and only the elements maturation has not yet reached. Prevalence is therefore cumulative rather than instantaneous: within a segment, whose elements enter their windows together, the interval between two prevalences is the interval between their exits, while between segments it also carries the difference in when the segments entered, about seven points against a range of seventy in the arm. Figure 1 is drawn from Willert and Henkel (1969), 169 children seen consecutively at the Heidelberg orthopaedic clinic between 1958 and 1962. Its thirteen limbs with a triphalangeal thumb are excluded from the denominator, a duplication not being a reduction; on the reduced denominator digit 1 comes to exactly 100%, the value the coded cohort reaches independently. Elements drawn with a solid line are counted directly, every limb of a sub-type carrying the same state of digit 1, the radius, the humerus and the ulna as stated in that sub-type’s own text, with Abb. 2 giving each sub-type’s size; those drawn dashed are derived from the mean finger number Abb. 22 publishes for each sub-type, distributed by the radial-to-ulnar rule the authors state on p. 53 in a split that reproduces the published mean exactly. The unpermitted combination reduces the humerus, the radius and all five rays while the ulna remains — digit 5 lost with the ulna standing between it and the humerus, the one arrangement the proximo-distal rule forbids. All three instances come from one investigator’s alizarin-stained fetal material, two in bonnet monkeys (Hendrickx and Newman, 1973) and one in a green monkey (Hendrickx and Sawyer, 1978); the 76 macaque forelimbs of the radiographic series show none. Alizarin red stains mineralised bone only, the carpals were unossified at examination by those authors’ own account, the digits ossify after the long bones, and the series mixes fetuses examined at 70 and at 100 days without stating which entry is which — so a ray recorded as absent there may be a ray not yet ossified. Hendrickx and Newman rank their own material the same way the order does: on the right side the ulna was unaffected and the fifth ray affected only rarely. A co-reduction was scored as consistent with one exposure when every element that leaves its sensitive period later than the deepest element reduced is itself reduced, with no allowance made for elements whose prevalences coincide. Scored this way, a single shared sensitive window accounts for 140 of the 142 human limbs, and their reduced elements form an exact prefix of the antero-posterior order (Supplementary file 3); all 14 observed co-reductions lay within the 35 antero-posterior-permitted patterns (Table 1). Four further limbs are scored elementwise but reach no combination because an element is unassessed; counting every unassessed element in them as spared, the least favourable reading, none takes a combination the two rules forbid. The truncation category is rare in the arm and common in the leg: seven upper limbs of six individuals meet it, five human children and one macaque, against 25 lower limbs, seven human from four children and eighteen primate from nine animals. Both limbs the shared window fails to account for belong to one of those six, and both fall inside the category; every one of the 136 limbs outside it is accounted for. The cohort’s only shortfall under this scoring was the left hand of Lenz and Knapp 600909, and it was removed not by a judgement about the phenotype but by a documentary correction. The workbook carried the verbatim “Klumphand links, Radiusaplasie, 3 Finger links”; the source (Abb. 6b, p. 1240) reads “Klumphände, Radiusaplasie, links 3. Finger, rechts Syndaktilie zwischen Zeige-und Mittelfinger”. Lenz and Knapp spell cardinal finger counts as words throughout — “beiderseits nur drei Finger”, “Links sechs Finger”, “Rechts drei Finger ohne Daumen” — and write a numeral with a full stop for an ordinal, as in “3. Finger und 5. Zehe beiderseits doppelt”. “links 3. Finger” is therefore the left third finger and gives no count for that hand, so the number and the identity of its reduced rays are both unstated, the autopod is unassessable, and the limb keeps every per-element prevalence while leaving the combination cohort. We report this explicitly because the correction favours the present hypothesis in every direction it touches: this limb was the only one outside the reduction sequence, the only shared-window shortfall, and one of only two placing digit 2 before the humerus. No other limb was re-examined on this occasion and no other limb moved.

### The model, the table and the figure

The two rules are stated as constraints on sets and applied by enumeration. Eight elements give 255 non-empty co-reductions. The A–P reduction rule forbids any set holding an element while a more anterior element of the same segment is absent from it, which leaves 35; the P–D reduction rule forbids any set holding a more distal element of the postaxial column while a more proximal one is absent, which leaves 25. Table 1 lists those 25 against the 14 the two cohorts show, and Supplementary file 2 the 12 that no species exhibits. No parameter is fitted at any point; the enumeration is exhaustive and the rules either admit a set or they do not.

The model is the two rules together with the prevalences. The rules fix which orders are admissible and the prevalences say where each element falls within an admissible one; neither alone gives a sequence. Stated as an order, the rules assert seven relations in three groups, not two chains: within the autopod digit 5 leaves before digit 4, digit 4 before digit 3, digit 3 before digit 2 and digit 2 before digit 1; along the postaxial column digit 5 leaves before the ulna and the ulna before the humerus; and within the zeugopod the ulna leaves before the radius. Digit 5 and the ulna each belong to two of these, which is what interleaves the segments. Of the 40,320 orderings of eight elements, 70 satisfy all seven, so the rules alone reduce the space by a factor of 576. What they do not settle is the humerus against the radius: 35 of the 70 put the humerus first and 35 the radius, and prevalence decides it in both species. Maturation is not asserted to move from element to element in steps. It advances continuously on the two axes, and the elements are the points at which the advance can be observed, an element leaving its window when it resolves into a discrete bone. The observed sequence is therefore a reading of a continuous process at eight places, and its order is the order of the eight prevalences. Both cohorts give a reading that satisfies all seven: digit 5, digit 4, digit 3, ulna, digit 2, humerus, radius, digit 1 in the human material, and digit 5, ulna, digit 4, digit 3, humerus, digit 2, radius, digit 1 in the macaque. The two differ where the column crosses the digits — the ulna stands after three digits in man and after one in the macaque — and nowhere else.

Table 2 is the human order read as successive depths. Each row is the set of elements that have not yet left their windows after one further element has left, so row k holds the last 9 − k elements of the order and the rows are its eight prefixes read from the deepest. The counts are the human limbs whose co-reduction is exactly that set. Figure 1 places the same eight elements at their prevalences on a single vertical scale, which the nesting of the windows permits: an element with a higher prevalence has a window containing that of every element below it, so one axis carries both directions. Neither the table nor the figure introduces any quantity not already in the coded cohort.

### Software and reproducibility

All analyses were performed in Python with a single dependency, openpyxl; matplotlib is required only to draw Figure 1 and by no other script. A self-contained script (Source code 1, `reproduce_results.py`) regenerates from the Source Data spreadsheet the principal reported quantities — cohort counts, the A–P reduction rule, the per-element prevalences under both the strict A∪P and broad A∪P∪H definitions, the cross-species rank correlation, and the dose-calendar (timed-primate) analysis. Source code 3 (validate_all.py) reads the same file and nothing else, re-deriving every value above and asserting it against the numbers reported here, so the manuscript cannot drift from the cohort. Source code 2 (make_figure1.py) draws Figure 1 from Willert and Henkel’s published sub-type counts, tabulated in its header with the page of the monograph each is taken from; it reads no data file, and validate_all.py checks the panel it produces — its denominator, its ulna value, and that its order is the one the cohort returns — against the cohort itself.The script additionally supports per-limb verification (`reproduce_results.py –case <Row_ID>`), which prints the original source quotation, the raw element codes, and the derived reduction status for any individual limb, so that the path from each published source description to its code to every reported statistic is fully inspectable for the selected cases. A README documents how each command maps to each table, figure, and number. The package is deterministic and accompanies this preprint as Source data 1 and Source code 1–3.

## Acknowledgements

The author thanks the investigators whose clinical and experimental documentation across 1962–2025 made the cohort assembly possible.

## Additional information

### Ethics

This study is a secondary analysis of previously published human case series and previously published experimental non-human-primate series. No new human participants and no new animal experiments were involved, and all human material analysed is anonymous and already in the published record. Ethical approval and informed consent were therefore not required.

## Data availability

All data analysed in this study are included in the manuscript and its supporting files. The unified verbatim-coded cohort, together with every source sheet and the Source index listing all contributing publications, is provided as Source data 1 (`Cohen_2025_Source_Data.xlsx`). The three analysis scripts are provided as Source code 1–3 (`reproduce_results.py`, `make_figure1.py`, `validate_all.py`). A README documents how each command maps to each table, figure and number. The complete package — data, scripts and README — accompanies this manuscript as source-data files. All primary series analysed here were previously published and are cited in the reference list. Collins and Scott (2025) is the only source coded from images: every limb was read from the skeletal illustrations in their supplemental material. Two further series contribute limbs to the combination cohort and are not otherwise named here: Wilson and Gavan (1967), three macaque forelimbs, and Kamimura et al. (2024), nine human limbs imaged by computed tomography in survivors at 59 to 61 years — six decades of remodelling changes how an element looks but not whether it is there. Willert and Henkel (1969) and Kida (1987) were coded from the text and tables, with the published figures and radiographs used to check each case, except three Kida cases: two whose plates were never scanned and one whose two radiograph panels are unreadable. Their six limbs are held out of the analysed cohort pending legible images, because in this source the published plate revised the coding in nearly every patient whose film was read. Reinstating all six leaves the reduction order, the observed co-reductions, the zero A–P violations and the two shared-window shortfalls exactly as reported. The sheet “Source locators” of Source data 1 maps each of the 269 limbs drawn from these three sources to its figure, patient or animal in the original. All three are under publisher copyright and are not redistributed here; copies can be sent on request.

